# Analysis of companion cell and phloem metabolism using a transcriptome-guided model of Arabidopsis metabolism

**DOI:** 10.1101/2022.08.04.502830

**Authors:** Hilary Hunt, Nico Brueggen, Alexander Galle, Sandy Vanderauwera, Claus Frohberg, Alisdair R Fernie, Uwe Sonnewald, Lee J Sweetlove

## Abstract

Companion cells and sieve elements play an essential role in vascular plants and yet the details of the metabolism that underpins their function remain largely unknown. Here we construct a tissue-scale flux balance analysis (FBA) model to describe the metabolism of phloem loading in a mature Arabidopsis leaf. We explore the potential metabolic interactions between mesophyll cells, companion cells, and sieve elements based on current understanding of the physiology of phloem tissue and through the use of cell-type-specific transcriptome data as a weighting in our model. We find that companion cell chloroplasts likely play a very different role to mesophyll chloroplasts. Our model suggests that, rather than carbon capture, the most crucial function of companion cell chloroplasts is to provide photosynthetically-generated ATP to the cytosol. Additionally, our model predicts that the metabolites imported into the companion cell are not necessarily the same metabolites that are exported in phloem sap; phloem loading is more efficient if certain amino acids are synthesised in the phloem tissue. Surprisingly, in our model predictions the H^+^-PP_i_ase is the more important contributor than the H^+^ ATPase to the energisation of the companion cell plasma membrane.

## Introduction

The transport of sugars and other key nutrients such as amino acids from source leaves via the phloem to sink tissues is essential for the growth of plants (van Bel 2021). While xylem works through the capillary action of water and dissolved ions along lifeless conduits (Bollard 1960; Strasburger 1891), phloem tissue is composed of living, tube-like sieve elements (van Bel 2003). Although sieve elements are living cells they are generally considered almost metabolically inert whose primary purpose is as conductors of phloem sap (van Bel 2003; Esau 1939). Their symplasts are connected at each end by sieve plates which allow fluid flow and the passage of molecules such as sugars and amino acids (with size limits up to at least 67kDa (van Bel 2018; Stadler et al. 2005)). This flow is vulnerable to structural breaches, for example by herbivory and this triggers a blockage via callose (Barratt et al. 2011; van Bel 2006; Esau et al. 1953).

To facilitate their role as living pipes, sieve elements contain a stripped back cellular machinery. They are devoid of a nucleus, functioning photosynthetic chloroplasts (although they do contain other plastids capable of storing starch (Behnke 1991)), and vacuoles (van Bel et al. 2002). In addition, their mitochondria are relatively rudimentary and, while metabolically active (Lee et al. 1971; McGivern 1957; Moninger et al. 1993), are thought unlikely to have the same capabilities as those in other cell types (Behnke et al. 1990; Cayla et al. 2015; Esau et al. 1953; Esau and Cronshaw 1968). The roles of these organelles are taken up by adjacent companion cells. Companion cells are symplastically linked to adjacent sieve elements so that metabolites can flow freely between them (Goodwin 1983; Stadler et al. 2005). Necessary mRNA transcripts are produced in companion cells for use in sieve elements (Ayre et al. 2003; van Bel 1996; Cayla et al. 2015) as well as small (<67 kDa) proteins (Stadler et al. 2005).

It is well established that osmotic currents drive the flow of sap through phloem (De Schepper et al. 2013; Knoblauch et al. 2016; Münch 1930). This means that the energy cost of phloem transport is primarily in the maintenance of the concentration gradient driving these osmotic currents. In the source leaves of *Arabidopsis* and similar herbaceous angiosperms, companion cells play an important role in actively importing photoassimilates produced in the mesophyll into the phloem tissue to maintain this gradient (Otero and Helariutta 2017). Sugars and amino acids produced in the leaf are passively exported from mesophyll cells into the apoplast. The bioenergetic force behind the loading of sugars and amino acids from the apoplast into companion cells is generated by proton gradients at the companion cell plasma membrane (Taiz 2010). The proton gradient across the plasma membrane is maintained by actively pumping protons from the companion cell cytosol to the apoplast. This allows companion cells to use proton-coupled plasma membrane symporters to import the photoassimilates in the apoplast against their concentration gradients but down the proton concentration gradient (Haritatos et al. 2000; Taiz 2010).

The proton motive force (PMF) is created and maintained by proton pumping ATPases (H^+^-ATPases) and proton pumping pyrophosphatases (H^+^-PP_i_ases) (Hafke et al. 2005; Langhans et al. 2001; VanBel 1995; Zhen et al. 1997). Both have been localised to companion cell membranes (DeWitt and Sussman 1995; Paez-Valencia et al. 2011; Pizzio et al. 2015). These molecular pumps can harness the energy released from hydrolysing ATP or PP_i_ to pump protons out of the cell against the concentration gradient. It is unclear to what extent the generation of the companion cell membrane PMF is split between these two pumps. Companion cell-specific overexpression of H^+^-PP_i_ase increases plant growth (Khadilkar et al. 2016) and knockdown produces dwarf plants (Primo et al. 2019). Similarly, hyperactivation of H^+^-ATPase may also increase plant growth under low pH conditions (Robertson et al. 2004) while knockout of the companion cell-specific H^+^-ATPase in tobacco also results in dwarf plants (Zhao et al. 2000). However, in Arabidopsis, homozygous knockout of the companion cell-specific H^+^-ATPase was not achieved due to its potential role in pollen development (Robertson et al. 2004). These results suggest that the capacity of the proton pumps is potentially a limiting factor for phloem transport and plant growth. They also demonstrate some redundancy in the role of the two proton pumps in maintenance of the companion cell PMF, perhaps making it more robust to internal substrate availability, i.e., ATP or PP_i_.

There has been speculation that, given the presence of two pumps and the thermodynamics involved, the plasmalemmal H^+^-PP_i_ases may actually be working in reverse – using the PMF to synthesise PP_i_ (Davies 1997; Gaxiola et al. 2016; Khadilkar et al. 2016; Rocha Façanha and de Meis 1998; Scholz-Starke et al. 2019). One reason put forward for generation of PPi in this way is that it could be used as a metabolic energy source to reduce the ATP cost involved in sucrose degradation (Gaxiola et al. 2012; Gutiérrez-Luna et al. 2018; Igamberdiev and Kleczkowski 2021; Sonnewald 1992) allowing companion cells to maximise ATP generation through respiration of sucrose. The high ATP consumption required to maintain companion cell PMF and the high sucrose concentrations in companion cells underpin the arguments for this alternative sucrose degradation pathway. Most persuasively, SUC1 transcript abundance, a sucrose-proton symporter, was found to increase in *Arabidopsis* plants overexpressing H^+^-PP_i_ase (Gaxiola et al. 2012; Gonzalez et al. 2010).

Because PP_i_ is generated from a variety of biosynthetic reactions including synthesis of ribonucleic acids and protein, a full understanding of the respective contributions of the plasma membrane H^+^-ATPase and H^+^-PP_i_ase pumps to companion cell energetics can only be reached after a comprehensive analysis of fluxes throughout the companion cell metabolic network. Recently, two Arabidopsis leaf cell-specific transcriptome studies were published that were able to provide some clues as to metabolic function in the companion cell (Kim et al. 2021; You et al. 2019). However, it is difficult to say anything definitive about enzyme concentration, let alone reaction flux, solely from transcriptome data (Lan et al. 2012; Peng et al. 2018; Schwender et al. 2014). We therefore decided to integrate the published Arabidopsis cell specific transcriptome data with a new leaf-scale flux-balance analysis (FBA) model of central metabolism that incorporates the metabolism of the mesophyll, companion cells and phloem sieve elements, the goal being to resolve uncertainties about the metabolism of companion cells and sieve elements. Previous leaf FBA models with biomass accumulation or phloem output as their objective function have been able to give real insight into cell metabolism despite their lack of identifiability (Cheung et al. 2014, 2015; Nikoloski et al. 2015; Shameer et al. 2019; Sweetlove and Ratcliffe 2011). Integrating transcriptome data into such models using carefully-considered algorithms that look at transcriptome data across all metabolic pathways covered by the model not only reduces the potential solution space of the model but can act as a weighting so that enzymes with high transcript abundance are more likely to carry high flux in the chosen model solution (Robaina Estévez and Nikoloski 2014; Robaina-Estévez et al. 2017; Scheunemann et al. 2018). Here we describe the results of detailed analysis of this model and discuss the implications of our findings for the efficiency and metabolic underpinnings of phloem loading.

## Results

### A tissue-scale model of metabolism of an Arabidopsis source leaf incorporating phloem loading

To develop a tissue scale model of phloem metabolism, we started with a diel FBA model of central plant cell metabolism in the leaf (Shameer et al. 2018, 2019) and adapted it to describe the cells involved in phloem loading. We determined the most necessary of these to be mesophyll cells, companion cells, and sieve elements – the cells most implicated in the synthesis, loading, and long distance transport of the photoassimilate component of phloem sap respectively (van Bel 1996; Haritatos et al. 2000). We included sieve elements in the leaf and sieve elements in the petiole as two distinct cell types in an effort to capture the continuous cost of sucrose retrieval and cell maintenance away from the energy-rich source leaf.

A schematic summary of the model is shown in Figure 1. Each cell type was modelled as a distinct diel FBA cell – composed of two metabolic models one each for the light phase and the dark phase with these two phase models connected by accumulation reactions that mimic the build-up of starch, sucrose, nitrate, malate, and amino acids during either the light or dark phase for use in the opposite phase (Cheung et al. 2014). The difference between the light phase and dark phase models is the flux of incident photons which is only present in the light phase model. Such models were constructed for each of the four cell types being considered. The light phase cell models were connected via a light phase apoplast compartment and the dark phase cells similarly connected via a dark phase apoplast compartment. Companion cells and sieve elements were connected via the symplast using no-cost ‘linker’ reactions that allowed free exchange of the majority of model metabolites. We excluded large molecules from this symplastic transport, such as heteroglycans, polysaccharides (e.g. cellulose and starch), and lipids as well as superoxide, protons, and iron ions from these transport reactions based on the size limit in companion cell-sieve element plasmodesmata and the unrealistically high transport fluxes we initially saw transporting charge between cells. Nucleotides were only allowed to travel from companion cells to sieve elements but not from sieve elements to companion cells to prevent similarly unlikely energy transport between cells. Without these exclusions, the model transported substantial quantities of ATP from companion cells to sieve elements via energy rich lipids or directly as ATP or equivalent nucleotide triphosphates. The petiolar sieve elements component of the model included a phloem export reaction that carried photoassimilates to the bulk phloem in the ratios measured by Wilkinson and Douglas (2003). The dark phase phloem output flux was constrained to a third of the light phase phloem output flux. Upper-bound constraints were placed on all fluxes in each phloem cell type to reflect the relative quantity of these cells relative to mesophyll cells in a mature Arabidopsis leaf (see Methods).

**Figure 1.**
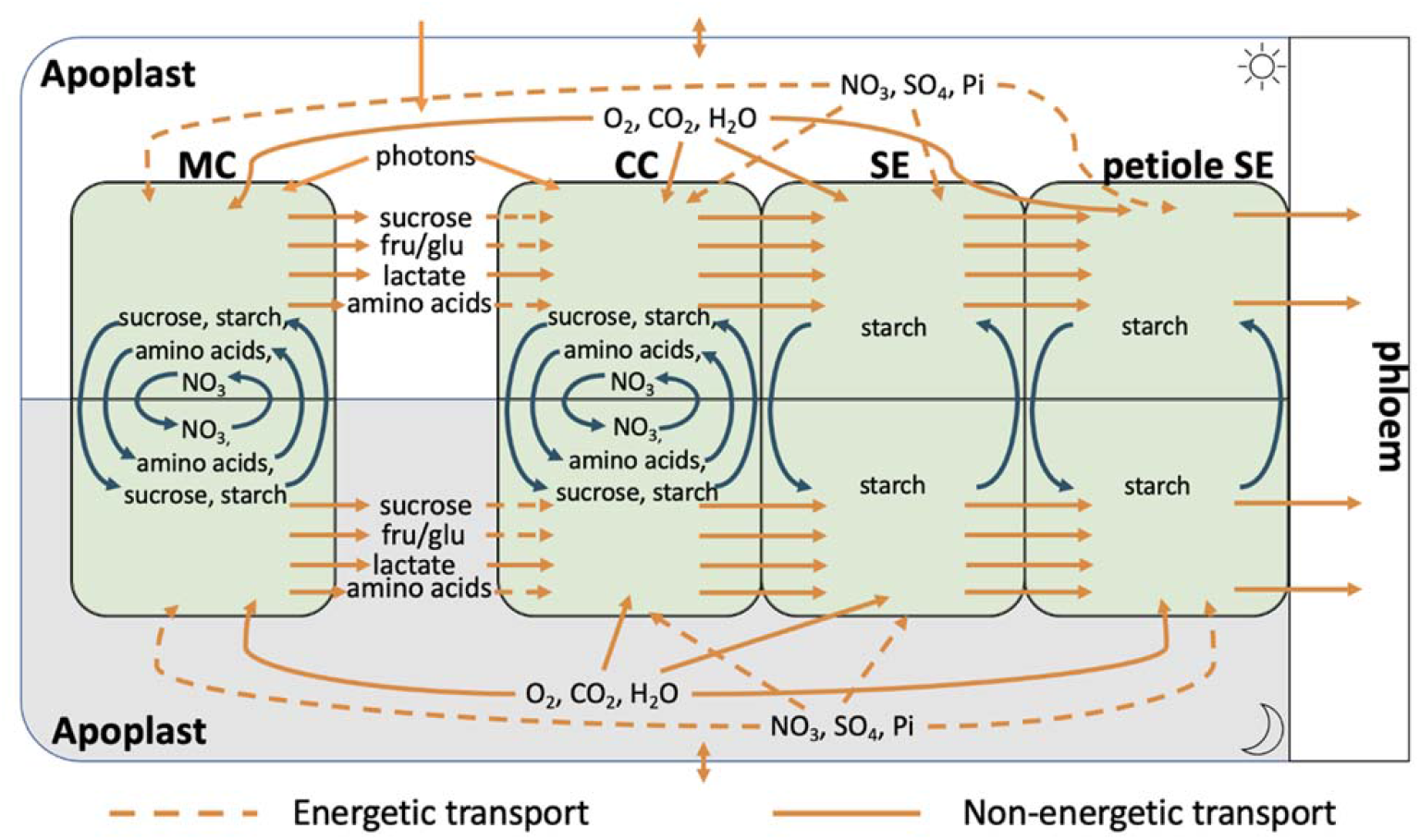
Schematic of the diel-multi-cell-FBA model of leaf tissue showing the inputs, outputs and exchanges of metabolites, gases, and minerals between the four cell types considered. Each cell contains a core stoichiometric model of central metabolism and fluxes are scaled to account for the different proportions of each cell type in the leaf. Each cell model is further replicated to account for the light phase (above) and the dark phase (below) with certain metabolites and minerals allowed to be passed between the phases as indicated by blue arrows. Since sieve elements do not have vacuoles, only starch is allowed to accumulate in them. The cells are all connected via the apoplast (this is also split into light phase apoplast and dark phase apoplast). However, only the companion cell, sieve element and petiole sieve element cells are symplastically connected; sugars and amino acids must transit through the apoplast in order to move between the mesophyll and companion cells. CC, companion cell; MC, mesophyll cell; SE, sieve element (in the leaf); petiole SE, sieve element in the petiole, fru = fructose, glu = glucose

Our model was further constrained by limiting the ribulose bisphosphate carboxylase-oxygenase flux in mesophyll cells so that the total model carbon assimilation rate matched the rate measured by Donahue et al. 1997. Cell maintenance costs were estimated for mesophyll cells based on the incident light as implemented by Töpfer et al., 2020. Maintenance costs for other cell types were then determined based on comparative cell numbers. Protein and mRNA turnover costs were also estimated based on experimental measurements (Li et al. 2017; Piques et al. 2009; Sorenson et al. 2018; Szabo et al. 2020) and imposed as model constraints. A sucrose leak and a sucrose retrieval flux were added to petiolar sieve elements based on data from Gould et al. 2012. All vacuole metabolic reactions were removed from sieve elements, including vacuolar metabolite accumulation. Due to the observed presence of starch-containing plastids (Behnke 1991), starch accumulation was allowed in sieve element plastids although the reaction carried no flux in our model solutions. Transport of amino acids through the apoplast was constrained to only flow in the direction of mesophyll to companion cell.

We used the cell type-specific transcriptome data collected by Kim *et al*., 2021 from 6 week old *Arabidopsis* plants to weight model reactions using a variation of the RegrEx algorithm (Robaina Estévez and Nikoloski 2014). The original RegrEx algorithm was designed to be applied to a model of a single cell type and determined the relative weighting of transcriptome data and the total sum of fluxes through regression. It found the model solution that best fit the available data while minimising the total sum of fluxes in the model to minimise the cost of enzyme turnover. RegrEx has since been used to model multicellular tissues (Robaina-Estévez et al. 2017; Scheunemann et al. 2018) but these studies relied solely on transcript abundance datasets to distinguish the cell types in the model. Given the substantial known differences between the cell types we modelled and the comparative, cell-specific transcriptome data available (Kim et al. 2021), we decided to use our more tailored FBA model. Combining evidence of cell structure and interconnectedness further reduced the degrees of freedom within our model and produced a more guided solution than RegrEx and a genome scale model alone, eliminating the bulk of physiologically impossible solutions from the outset.

For the purposes of our study, the RegrEx algorithm was modified by scaling transcriptome data to match cell ratios and the weighting of transcriptome data versus sum of fluxes. In the case of companion cell and sieve element reactions where companion cells synthesise the transcripts for both cell types, the objective was to minimise the difference between the sum of fluxes in companion cells and both sieve elements and their corresponding transcript abundance in companion cells. Previous analysis of transcriptome and metabolic flux in plants suggested an overall correlation between transcripts and flux of 32% (Schwender et al, 2014) and hence, the transcriptome weighted model solution was constrained so that the difference between its phloem output and that of a parsimonious solution (see Methods for more detail) was no greater than 32%. This meant including a constraint on the minimum phloem output to ensure phloem loading activity. The model was then solved as a single optimisation problem with the objective of minimising the difference between modelled reaction flux and scaled transcriptome abundance within these constraints. To ensure the robustness of model predictions, we also performed flux variability analysis (FVA) on the transcriptome-weighted model to determine the ranges the flux through each reaction could take while maintaining the shortest distance from the transcriptome data. Our model suggests the likely distributions of ATP generation and consumption, as well as likely metabolic activity in companion cells and sieve elements. For full details see the Methods section.

The addition of phloem cells to the diel leaf model does not have a pronounced effect on the metabolic behaviour of mesophyll cells with flux distributions (see Supplemental Table ST1; Supplemental Fig. S2) broadly similar to those reported previously (Cheung et al. 2014, 2015; Shameer et al. 2019). These included the requirement for substantial mitochondrial ATP synthesis in the mesophyll in the light phase as well as export of reducing equivalents from the chloroplast via the malate valve (Shameer et al. 2019). The broad patterns of metabolite accumulation in day or night were also similar to previous reports (Cheung et al. 2013, 2014, 2015).

### Companion cell photosynthesis functions in the model to generate reducing power and ATP but with no carbon fixation

One of the most notable features of the model solution was the unusual metabolism in companion cell chloroplasts. We modelled companion cell chloroplasts as metabolically identical to mesophyll cell chloroplasts, i.e. the same reactions were allowed. However, we imposed a constraint on the maximum rate of photosynthesis in companion cell chloroplasts reflecting the comparative light capturing capacity of mesophyll and companion cells due to the different proportions of each cell per unit leaf and the significantly reduced total chloroplast volume in companion cells (Cayla et al. 2015; Paramonova et al. 2002). These amount to restrictions on companion cell photon capture (Photon_ep) to 20% of the upper bound constraining mesophyll cell photon capture on a per cell basis (Supplemental Table ST1).

With these constraints, companion cell chloroplasts were predicted by the model to still generate approximately half the amount of ATP as mesophyll cell chloroplasts on a per cell basis. This constituted 58% of the total light phase ATP needs of the companion cell (Supplemental Figure S3). This was achieved because a greater proportion of the light absorbed by companion cells was used for photochemistry in the model. This suggests a lower proportion of non-photochemical quenching occurs in companion cell chloroplasts compared to the average leaf chloroplast which is known to be substantial at the light intensity used in our model (Li et al. 2009; Müller et al. 2001). It also implies that the companion cell could potentially have an energy-limited, rather than carbon-limited metabolism.

The majority of companion cell photosynthesis-derived energy (51%) was predicted to be used in amino acid biosynthesis pathways in the chloroplast, while the rest was exported to the cytosol using either the glyceraldehyde 3-phosphate – 3-phosphoglycerate shuttle or the pyruvate-phospho*enol*pyruvate shuttle (Figure 2) to maintain the plasma membrane proton gradient necessary to continue sucrose and amino acid import. No carbon fixation by RuBisCO was predicted to occur in companion cell chloroplasts in the model. In contrast, in mesophyll chloroplasts, the model predicted that 80% of photosynthetically-generated ATP is used to fix carbon leaving the other 20% for use in other cellular processes.

**Figure 2.**
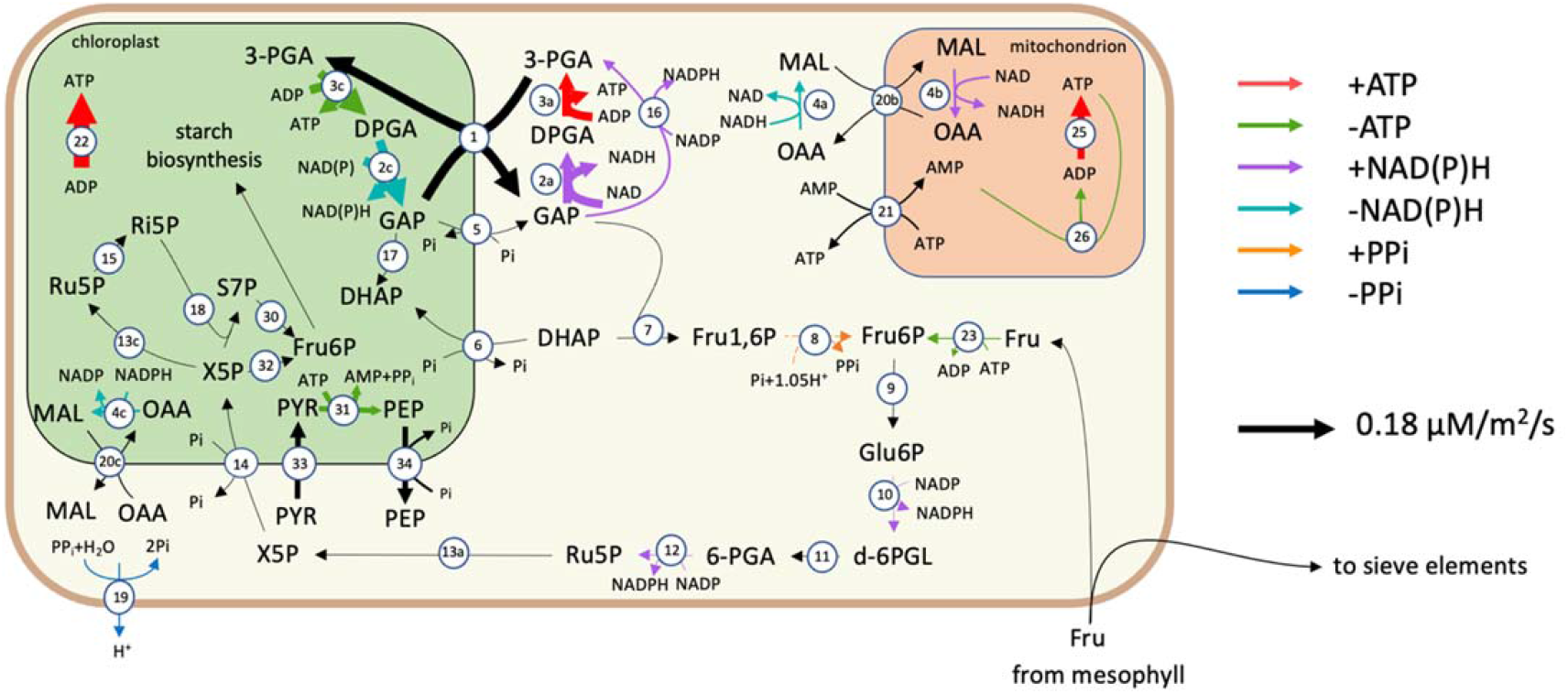
A flux map of the principal reactions involved in companion cells during the light phase. Arrow thickness indicates reaction flux value. Red/green arrows indicate ATP production/consumption, purple/teal arrows indicate NAD(P)H production/consumption, orange/blue arrows indicate PP_i_ production/consumption. The reactions shown are 1: GAP-3-PGA shuttle, 2: (NADP) GAP dehydrogenase, 3: 3-PGA kinase, 4: Malate dehydrogenase, 5: GAP-Pi shuttle 6: DHAP-Pi shuttle, 7: FBP aldolase, 8: FBP phosphotransferase, 9: Glu6P isomerase, 10: Glu6P dehydrogenase, 11: 6-phosphogluconolactonase, 12: Gluconate-6-phosphate dehydrogenase, 13: Ru5P epimerase, 14: X5P-phosphate shuttle, 15: Ribose-5-phosphate isomerase, 16: 3-PGA dehydrogenase, 17: Triosephosphate isomerase, 18: Transketolase, 19: H^+^-pyrophosphatase, 20: Malate-oxaloacetic acid shuttle, 21: ATP-AMP shuttle, 22: Plastidial ATP synthase, 23: Fructose kinase, 25: Mitochondrial ATP synthase, 26: Adenylate kinase, 30: Transaldolase, 31: Pyruvate orthophosphate dikinase, 32: Transketolase, 33: Pyruvate channel, 34: PEP channel, 35: Plastidial ATP synthase The metabolites shown are GAP: glyceraldehyde-3-phosphate, DPGA: 3-phospho-d-glyceroyl phosphate, 3-PGA: 3-phosphoglycerate, DHAP: Dihydroxy acetone phosphate, MAL: malate, OAA: oxaloacetic acid, Glu1P: glucose-1-phosphate, Glu6P: glucose-6-phosphate, d-6PGL: 6-phosphoglucono-∂-lactone, 6-PGA: 6-phosphogluconate, Ru5P: ribulose-5-phosphate, X5P: xylulose-5-phosphate, Ri5P: ribose-5-phosphate

A study of chloroplast structure (Paramonova et al. 2002) showed that companion cell chloroplasts have less pronounced grana than mesophyll cell chloroplasts. This could imply reduced or non-existent function of photosystem II (Dekker et al. 2002; Dekker and Boekema 2005) in these cells. We analysed the effect that removal of companion cell photosystem II had on the model solution and found that, while this did reduce the total amount of ATP available in companion cells (by 35%), companion cell photosynthesis still provided a substantial proportion of ATP used in the companion cell (56% versus 58% when photosystem II was present) and there was still no carbon fixation (Figure 3A, B). The lack of photosystem II meant that thylakoid plastoquinone needed to be reduced via the ferredoxin-plastoquinone reductase pathway and this consumed all of the electrons released by photosystem I eliminating NADPH synthesis. The NAD(P)H requirements of the Calvin-Benson cycle were fulfilled by importing reducing power from the cytosol via the malate-oxaloacetic acid shuttle and as a biproduct of the serine and histidine biosynthesis pathways. The lack of reducing power in companion cells greatly reduced mitochondrial ATP synthase activity as well (by 30.6%) (Supplemental Table ST1). However, the companion cell chloroplast was still capable of generating enough ATP to minimise the need for catabolism of metabolites imported into the companion cell (Supplemental Table ST1).

**Figure 3.**
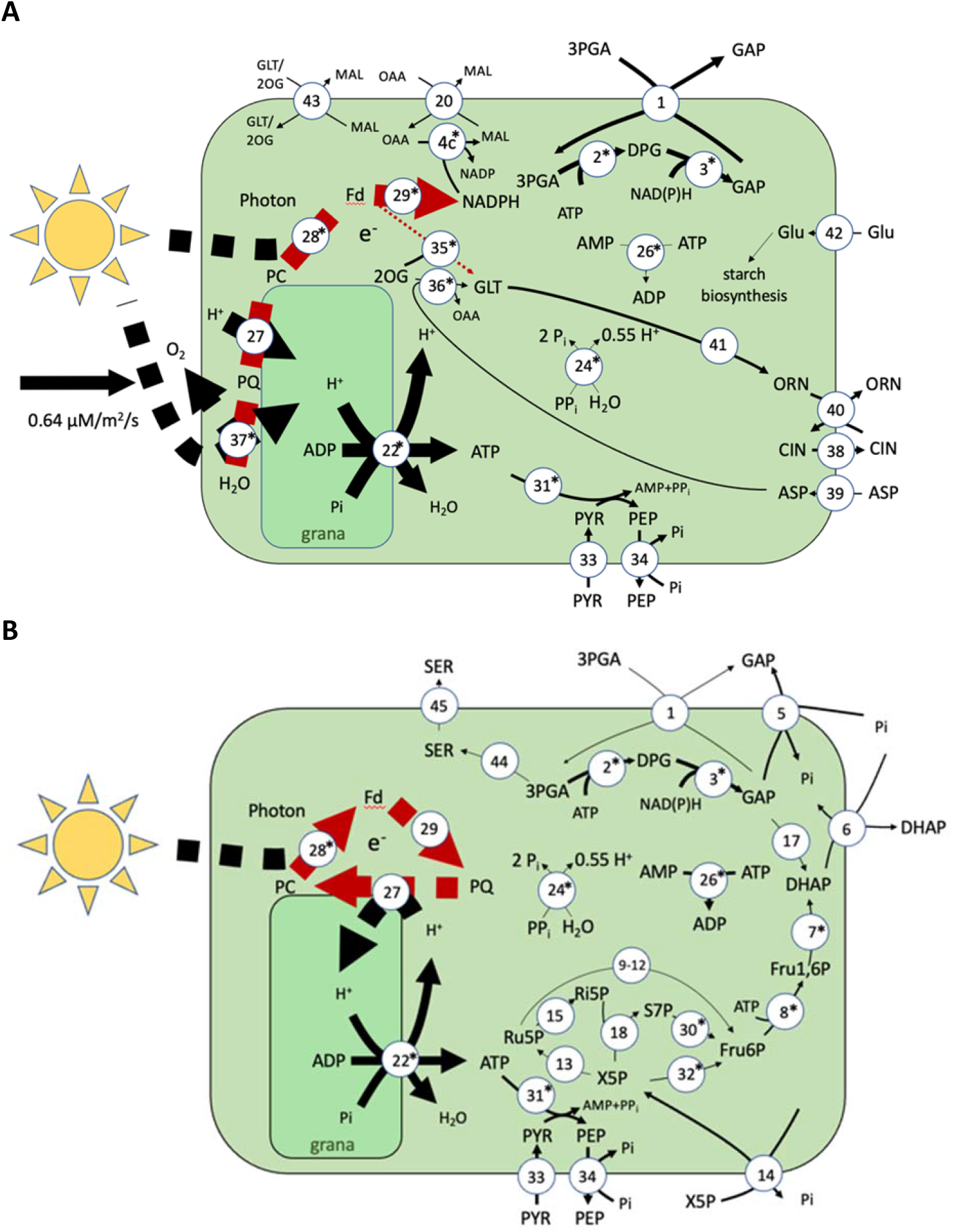
Energy metabolism in companion cell chloroplasts. (A) A flux map of the transcript-guided model solution in the light phase companion cell chloroplast. The dashed, maroon arrows indicate electron transport rather than mass flow. (B) A flux map of the transcript-guided model solution in the light phase companion cell chloroplast when PSII is disabled. Asterisks next to reaction numbers indicate a lower transcript abundance in companion cells when compared with mesophyll cells. Reactions with flux less than 0.005 µM/m^2^/s were excluded. The thickness of the lines of each solid arrow (excluding electron transport reactions) is scaled linearly to flux. The scale is shown on the left hand side of each diagram. The reactions shown are 1: GAP-3-PGA shuttle, 2: (NADP) GAP dehydrogenase, 3: 3-PGA kinase, 4: NADP^+^ Malate dehydrogenase 5: GAP-Pi shuttle 6: DHAP-Pi shuttle, 7: FBP aldolase, 8: FBP phosphotransferase, 9: Glu6P isomerase, 10: Glu6P dehydrogenase, 11: 6-phosphogluconolactonase, 12: Gluconate-6-phosphate dehydrogenase, 13: Ru5P epimerase, 14: X5P-phosphate shuttle, 15: Ri5P isomerase, 16: Transketolase, 17: Triosephosphate isomerase, 18: GDP kinase, 19: H^+^-pyrophosphatase, 20: Malate-oxaloacetic acid shuttle, 21: ATP-AMP shuttle, 22: Plastidial ATP synthase, 23: GDP-glucose pyrophosphorylase, 24: Inorganic pyrophosphate, 26: Adenylate kinase, 27: Plastoquinol-plastocyanin reductase, 28: PSI, 29: Ferredoxin-plastoquinone reductase, 30: Transaldolase, 31: Pyruvate orthophosphate dikinase, 32: Transketolase, 33: Pyruvate channel, 34: PEP channel, 35: Glutamate synthase ferredoxin reaction, 36: Aspartate aminotransferase, 37: PSII, 38: Citrulline channel, 39: Aspartate channel, 40: Ornithine-citrulline shuttle, 41: Ornithine synthesis pathway (from glutamate), 42: Glucose channel, 43: Malate-glutamate and malate-2-oxoglutarate shuttles, 44: Serine biosynthesis pathway, 45: Serine channel The metabolites shown are GAP: glyceraldehyde-3-phosphate, DPGA: 3-phospho-d-glyceroyl phosphate, 3-PGA: 3-phosphoglycerate, DHAP: dihydroxy acetone phosphate, MAL: malate, OAA: oxaloacetic acid, Glu1P: glucose-1-phosphate, GDP-Glu: GDP-glucose, Glu6P: glucose-6-phosphate, d-6PGL: 6-phosphoglucono-∂-lactone, 6-PGA: 6-phosphogluconate, Ru5P: ribulose-5-phosphate, X5P: xylulose-5-phosphate, Ri5P: ribose-5-phosphate, Fd: ferredoxin, PC: plastocyanin, PQ: plastoquinol, Glu: glucose, GLT: glutamate, GLN: glutamine, PYR: pyruvate, PEP: phosphoenol pyruvate, CLN: citrulline, ORN: ornithine, SER: serine

### The transcriptome-weighted model indicates that H^+^-PP_i_ase is the dominant plasma membrane proton pump in companion cells

There is evidence of both H^+^-ATPase (AHA) and H^+^-PP_i_ase (AVP1) pumps on the plasma membrane of companion cells (Gaxiola et al. 2016; Otero and Helariutta 2017; Paez-Valencia et al. 2011). Given the presence of both proteins, it could be that both pumps are in use and important in maintaining the companion cell plasma membrane proton motive force (PMF), which is necessary for phloem loading. However, there is some debate about whether H^+^-PP_i_ase contributes to or consumes the PMF in companion cells (Davies 1997; Gaxiola et al. 2012; Langhans et al. 2001; Segami et al. 2018).

In our transcriptome-weighted model solution, only the H^+^-PP_i_ase carried flux. The transcript abundance for several pyrophosphate-synthesising reactions weights our model towards a solution in which sufficient PP_i_ is generated within the companion cells for H^+^-PP_i_ase to maintain the majority of the PMF required for uptake of sugars and amino acids. While there was no direct information for H^+^-PP_i_ase transcript abundance in the published dataset, pyrophosphate produced in protein/RNA turnover reactions and by pyrophosphate: fructose-6-phosphate phosphotransferase in the transcriptome-weighted model solution were sufficient for the H^+^-PP_i_ase pump to maintain the companion cell’s PMF with the modelled photoassimilate import. Additionally, with the PP_i_ produced by protein and mRNA turnover, FVA showed there was always a role for H^+^-PP_i_ase whereas the model solution can have zero H^+^-ATPase flux with the same phloem output and while being equally close to the transcriptome data.

### Uptake of metabolites from the apoplast into the companion cell

As expected, sucrose import is the largest flux of metabolites actively transported into the companion cell from the apoplast (Figure 4). Unexpectedly, there was a similarly high fructose import flux which was then used in glycolysis. The model predicted that the remaining energetic demands of the companion cells were largely met by companion cell photosynthesis and catabolism of imported amino acids. Approximately half of the imported sucrose (46%) was passed unchanged via the symplast into the phloem. The remaining portion (54%) was partially consumed, not within companion cells, but within the sieve elements to provide energy for cell maintenance and carbon skeletons for amino acid synthesis. – Approximately 70% of the modelled carbon fixed by mesophyll cell RuBisCO was exported in the phloem sap. Sucrose was catabolised in sieve elements either by sucrose synthase or invertase. This is consistent with evidence of sieve element specific sucrose synthase isoforms (Barratt et al. 2009; Yao et al. 2020) and the presence of both enzymes in the *N. tabacum* sieve element proteome (Liu et al. 2022).

**Figure 4.**
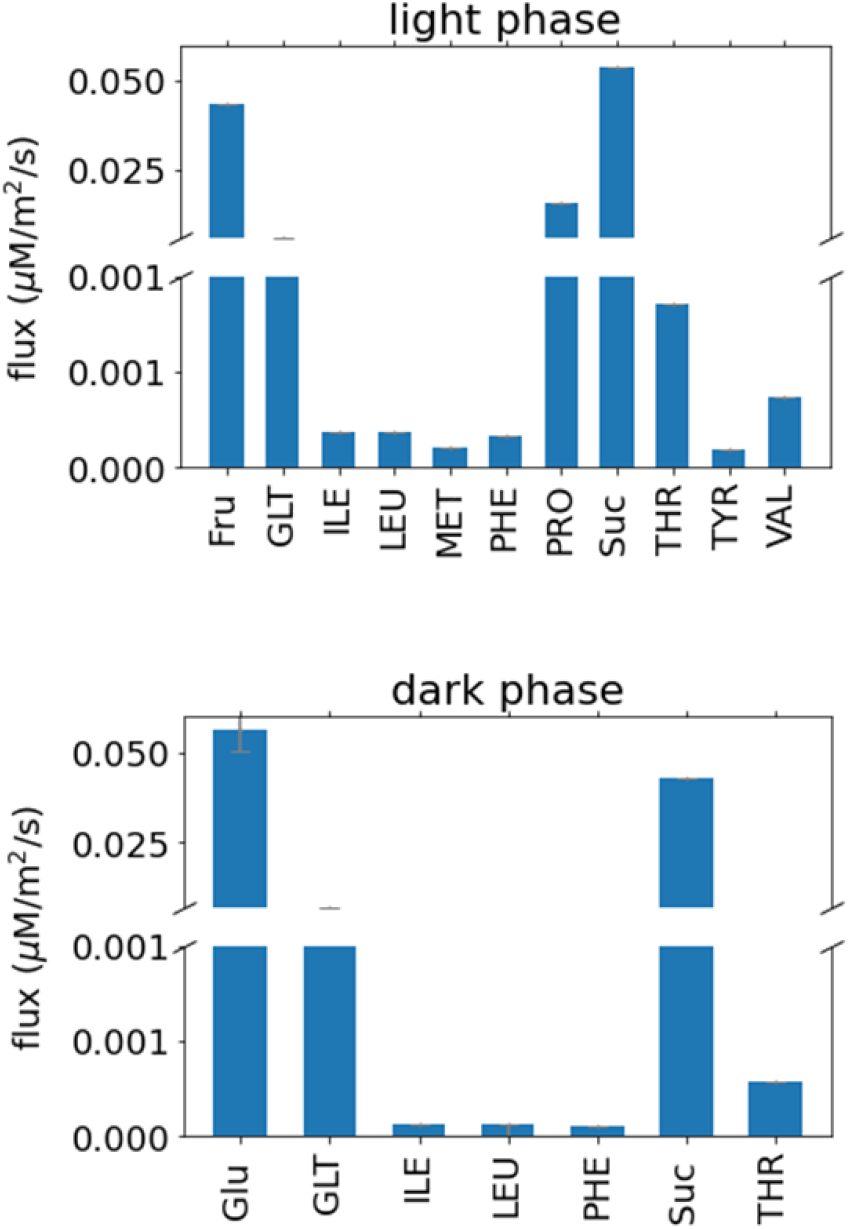
Predicted fluxes of metabolite transport via the apoplast between mesophyll and companion cells in the transcriptome weighted model solution. **Abbreviations are as in Figures 2, 3**.

Of the amino acids that were predicted to be taken up by the companion cell (Figure 4), most were transported unchanged through the connected symplast of companion cells and sieve elements, and into the bulk phloem. The exceptions to this were proline, glutamate and, in the light phase, aspartate and arginine. All four of these were used in the model to synthesise the remaining amino acids within phloem cells (Figure 5) and to provide additional ATP and reducing power (Supplemental Table ST1). While it is not exported in phloem sap, the model predicts that proline is also among the metabolites taken up by the companion cells from the apoplast. The extra reducing power obtained from converting proline into glutamate in phloem tissue via mitochondrial proline dehydrogenase and 1-pyrroline-5-carboxylate dehydrogenase allows the mesophyll cells to export fractionally more reducing power to the phloem. If this pathway was blocked in the model, proline is replaced with greater arginine and glutamate export to phloem cells and results in a small decrease in the maximum phloem output of the model.

**Figure 5.**
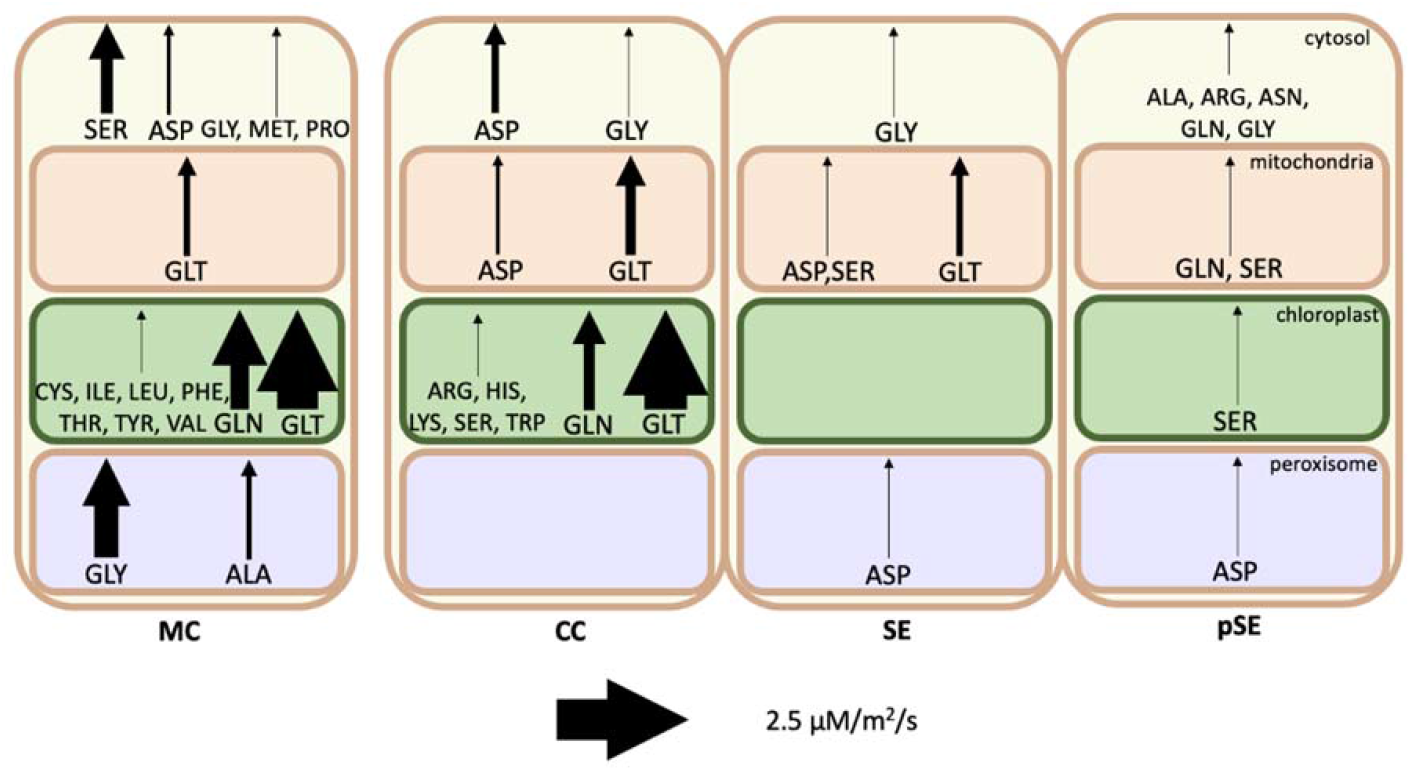
Amino acid synthesis by compartment in each cell in the light phase. Arrow thickness indicates magnitude of synthesis flux. Note that flux is scaled by cell ratios to better compare the activity between individual cells. MC: mesophyll cell, CC: companion cell, SE: sieve element, pSE: petiolar sieve element

### Amino acid biosynthesis could occur in phloem tissue

Many of the amino acids found in Arabidopsis phloem sap were imported into companion cells from the apoplast (and hence biosynthesised in the mesophyll) in our model solution. However, a not insignificant proportion were synthesised within the phloem tissue. Figure 5 depicts the relative amounts of amino acid biosynthesis occurring in each cell in our model solution during the light phase. In particular, our model suggests that arginine, asparagine, histidine, lysine, and tryptophan are more efficiently synthesised in phloem tissue from by-products of the oxidative pentose phosphate pathway and glycolysis than synthesised and transported from the mesophyll.

This result arises partly from the transcriptome weighting and partly from the optimisation objective of the model. Every cell in the model was predicted to carry flux through glycolysis and the oxidative pentose phosphate pathway to some extent to provide the ATP and NADPH necessary for cell maintenance. Of the five amino acids whose biosynthesis pathways branch off of these, our model predicted which would be most efficiently produced in the phloem, the main factor being the extent to which energy or reducing power were produced in their biosynthesis. This means that the histidine biosynthesis pathway which produces both NADH and pyrophosphate was active in companion cells in the model. Arginine, which is synthesised from aspartate, produces fumarate as a by-product and this fumarate was used by the model to produce ATP in phloem tissue (companion cells and petiole sieve elements) via the TCA cycle. Asparagine, which can be synthesised from aspartate and ATP while producing pyrophosphate, was also predicted to be synthesised in the petiole sieve elements where the pyrophosphate was used by the model in sucrose catabolism.

Surprisingly, tryptophan and lysine were also predicted to be synthesised in companion cells in our model. For tryptophan, the explanation is that several reactions in the indole synthesis pathway have higher transcript abundance in companion cells than in mesophyll cells and the synthesis of tryptophan consumes this indole. The prediction of biosynthesis of lysine in companion cells is surprising because it consumes ATP and reducing power, and reactions in its synthesis pathway generally had higher transcript abundance in mesophyll cells (Supplemental Table ST1).

In the dark phase when energy and reducing power are less abundant in mesophyll cells, and most other exported amino acids must be accumulated in the mesophyll to maintain phloem output, our model predicted that the cheapest of these same ‘low cost’ amino acids will still be actively biosynthesised in the phloem tissue at night. While amino acids produced in phloem tissue are produced in both the light and dark phase, other than glutamate, our model predicts that all amino acids produced in the mesophyll for phloem export must be produced during the light phase. They are then accumulated in the vacuoles for export in the dark phase.

Model solutions indicate that, apart from these and glycine, glutamine, alanine, aspartate, and serine which can be produced in either mesophyll or companion cells, the other amino acids are most efficiently produced in mesophyll cells.

### Adenylate kinase is necessary for model feasibility

Unexpectedly, we found that adenylate kinase (ATP+AMP -> 2ADP) flux is necessary for model feasibility and causes a net consumption of ATP in model solutions. Consistent with this, adenylate kinase mRNA has a relatively high abundance in the companion cell transcriptomes (Kim et al. 2021). Figure 6 shows the adenylate kinase fluxes and other reactions involving adenine nucleotides in the phloem cells. Despite the prediction of flux through the adenylate kinase reaction in all subcellular compartments in which it is present (mitochondrion, cytosol and chloroplast), there was no net adenine nucleotide transport between the cytosol and chloroplast in any cell type of the model.

**Figure 6.**
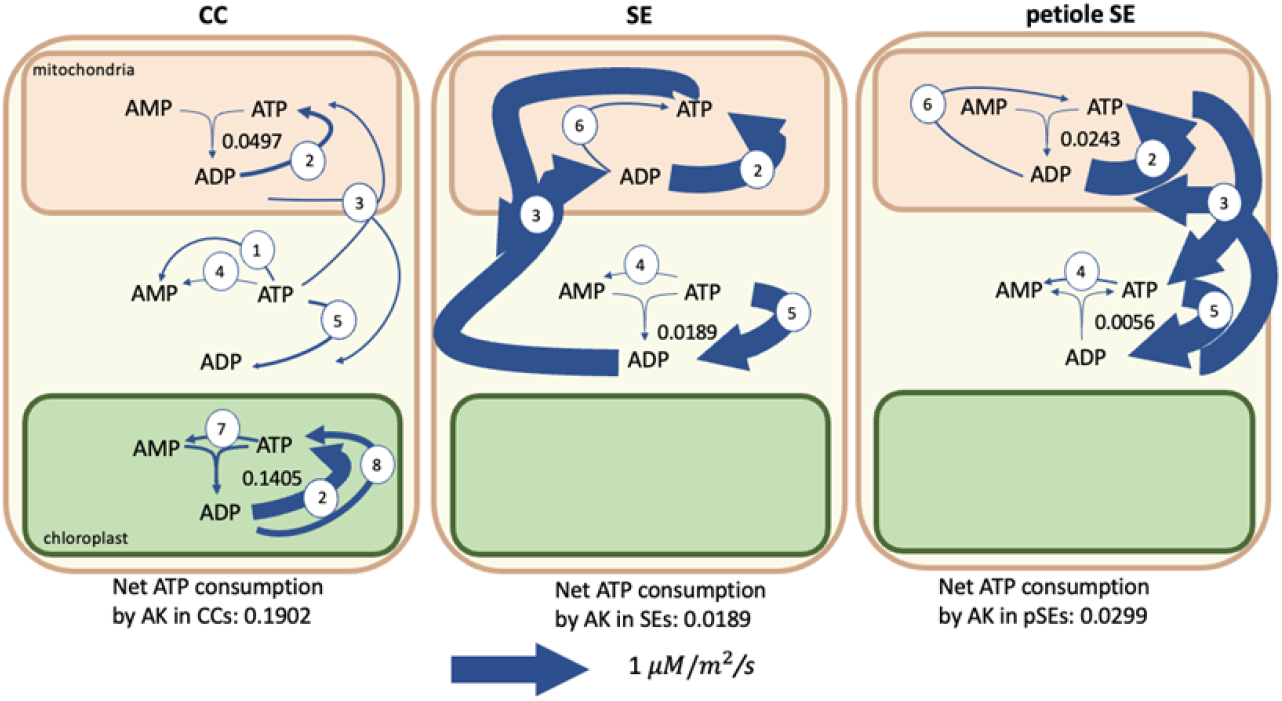
Flux map showing fluxes of adenylate kinases (AK) and related reactions in the transcriptome-weighted model solution. The net effect of adenylate kinase fluxes is a consumption of ATP. The reactions depicted other than adenylate kinase are 1: Pyruvate kinase, 2: ATP synthase, 3: ATP-ADP exchanger, 4: Protein/RNA turnover 6: Succinyl-coA synthetase, 7: Pyruvate orthophosphate dikinase, 8: Phosphoglycerate kinase and acetyl-glutamate kinase. Blue arrow thickness is scaled to flux value as indicated. Other than adenylate kinase, only fluxes > 0.05 µM/m^2^/s are shown. The total ATP consumption by adenylate kinase in phloem cells is 0.2390 µM/m^2^/s. CC = companion cell, SE = sieve element. The ranges on the adenylate kinase fluxes are: CC_m_: 0 – 0.1657, CC_c_: 0 – 0.1657, CC_p_: 0 – 0.1405, SE_m_: 0 – 0.1349, SE_c_: 0 – 0.1349, SE_p_: 0 – 0.1403, pSE_m_: 0 – 0.0299, pSE_c_: 0 – 0.0299, pSE_p_: 0 – 0 µM/m^2^/s ATP consumed.

Systematic *in silico* knockout of each adenylate kinase in the model showed that mitochondrial and cytosolic adenylate kinase activity in phloem cells together have the greatest effect on the rate at which the model was able to export sugars and amino acids to the phloem. Knockout of these two reactions in either companion cells or sieve elements resulted in an infeasible model. The primary function of these adenylate kinase isoforms in the model was to rephosphorylate AMP produced in other cellular processes. In the model solutions these were pyruvate orthophosphate dikinase for the chloroplast-cytosol PEP-Pi shuttle, Succinyl-coA biosynthesis in the TCA cycle, and protein/RNA turnover and resynthesis. Adenylate kinase activity in the mitochondrion allowed the model to balance the ratios of ATP, ADP, and AMP to match the organelle’s adenine nucleotide exchangers for optimal ADP phosphorylation via mitochondrial ATP synthase. This activity occurs in all phloem cells modelled - companion cells, sieve elements, and petiolar sieve elements.

### Differences between the transcriptome weighted model solution and an optimised parsimonious model solution

We compared the transcriptome weighted model solution with a parsimonious model solution that was not influenced by the transcriptome data at all. This latter solution was calculated to maximise phloem output and minimise model fluxes. The transcriptome weighted solution and the parsimonious solution have the same objectives and constraints but the transcriptome weighted solution is allowed a lower phloem output than the parsimonious solution so that it can also minimise the gap between the transcriptome data and model fluxes. This was allowed to give the model flexibility in case the transcriptome weighting led to less efficient metabolism overall. For example, the transcriptome data includes metabolic pathways such as cell wall and callose synthesis and fatty acid biosynthesis that would not be active in our mature-leaf model unless a specific constraint was added. There is also the possibility that some reactions carry a different flux *in vivo* than in our model because of thermodynamic and kinetic considerations. And it may also be the case that less efficient pathways are utilised in vivo due to evolutionary reasons (i.e. evolution does not necessarily optimise). These differences between the transcriptome-weighted model and an optimised model may point towards potential targets for engineering more efficient phloem loading with consequent positive impacts on crop performance (Braun et al. 2014; Gaxiola et al. 2016; Pazhamala et al. 2020).

The majority of the differences between the two model solutions can be linked to the difference in phloem output with most biosynthesis reactions running in the same cells, in the same ratios, but with reduced fluxes (Supplemental Table ST1). With the lower energy and carbon needs in phloem tissue in the parsimonious solution, the starch and sucrose accumulation in the companion cell chloroplasts and vacuoles that the transcriptome weighted solution predicts during the light phase for use in the dark phase is not necessary for optimal phloem output.

There were also differences in amino acid metabolism with the parsimonious solution only importing threonine, isoleucine, methionine and glutamate into the companion cells. The rest of the amino acids found in the phloem sap (and hence required in our model’s phloem output) were predicted to be synthesised within phloem cells (companion cells or sieve elements) in the parsimonious model. In general, there is an energetic or reducing power-based benefit in synthesising these amino acids in phloem cells allowing companion cells to be more efficient in importing only the most energy rich amino acids and synthesising the rest from them. Several, such as glycine, glutamate and glutamine are just as easily produced in phloem tissue as mesophyll cells based on stoichiometry. The differences are based on the transcriptome data and could indicate potential engineering targets to increase the efficiency of phloem transport.

Additionally, fructose was not imported into the parsimonious model’s companion cells. Companion cell chloroplasts in the parsimonious solution channelled all of the ATP and reducing power they produced into the GAP-3PGA export shuttle rather than the less efficient PEP-pyruvate shuttle. There was then enough ATP to satisfy cell maintenance costs, including protein and RNA turnover, and dephosphorylate fructose-1,6-bisphosphate to produce enough PP_I_ to import the required amino acids and sucrose. The less efficient PEP-pyruvate shuttle was used in the transcriptome weighted solution due to a high transcript abundance of pyruvate kinase in all cell types. While pyruvate kinase is important in the glycolytic pathway (Ambasht and Kayastha 2002), its primary use in our model was in facilitating the PEP-pyruvate conversion.

## Discussion

### Chloroplast activity in companion cells

Our model analysis suggests that given the energy demand of phloem loading, photosynthesis in companion cell chloroplasts is required for the most efficient system. The bulk of the ATP and reducing power generated in the companion cell chloroplast was exported to the cytosol via metabolite shuttles and there was no carbon fixation. Comparison of the structure of companion cell chloroplasts and mesophyll chloroplasts shows that those in companion cells have a thicker chloroplast peripheral reticulum with enlarged channels (Paramonova et al. 2002). This supports the likelihood of the high metabolite exchange between cytosol and chloroplast suggested in the model solution which transfers photosynthesis derived ATP (i.e. the GAP-3PGA shuttle) and could be indicative of the different role companion cell chloroplasts play. While there is no significant difference between abundance of RuBisCO transcripts in companion cells and mesophyll cells in the Kim et al 2021 dataset, several Calvin-Benson cycle enzymes and photosynthetic complexes do have significantly lower transcript abundance in companion cells compared with mesophyll cells (Figure 3A), including both photosystem I and II. In both the transcriptome weighted and parsimonious model solutions there was no flux through companion cell RuBisCO in contrast to the relatively high expression. To test this, companion-cell-specific knockdown of RuBisCO would be required, which is beyond the scope of the current study. Alternatively, if the Calvin cycle is operational in companion cell chloroplasts and consuming ATP to fix carbon then knockout of companion cell RuBisCO would be a promising knockout target for improved efficiency of phloem loading as it would allow more of the photosynthetically-generated ATP to be made available for this process.

It is also interesting to note that in model solutions where flux through companion cell photosystem II was allowed, companion cell mitochondria were predicted to produce large amounts of ATP (Figure 7) with the reducing power to facilitate this being derived from photosynthesis rather than from sucrose catabolism. The transfer of reducing power from chloroplast to mitochondrion is similar to that described by Bailleul et al., 2015 in diatoms but in our model was transferred via the chloroplast GAP-3PGA shuttle rather than via the malate valve as in diatoms. It is unclear what the respective capacities of these metabolite shuttles in companion cells are and so this may be an unrealistically efficient flux pattern and could contribute to an overestimation of the capacity of the companion to generate. Should this be the case, it is likely that catabolism of sucrose would need to play a role as a source of ATP production in companion cells in contrast to the model prediction that no sucrose is catabolised in the companion cells. In this context, it is interesting to note that an Arabidopsis mutant completely deficient in all isoforms of sucrose synthase (Fünfgeld et al. 2022), including those expressed in companion cells, was able to grow and develop normally. However, this does not mean that no companion cell sucrose catabolism was occurring because invertase activity could take over the role in sucrose catabolism as it appears to in potato sucrose synthase knockouts (Zrenner et al. 1995).

**Figure 7.**
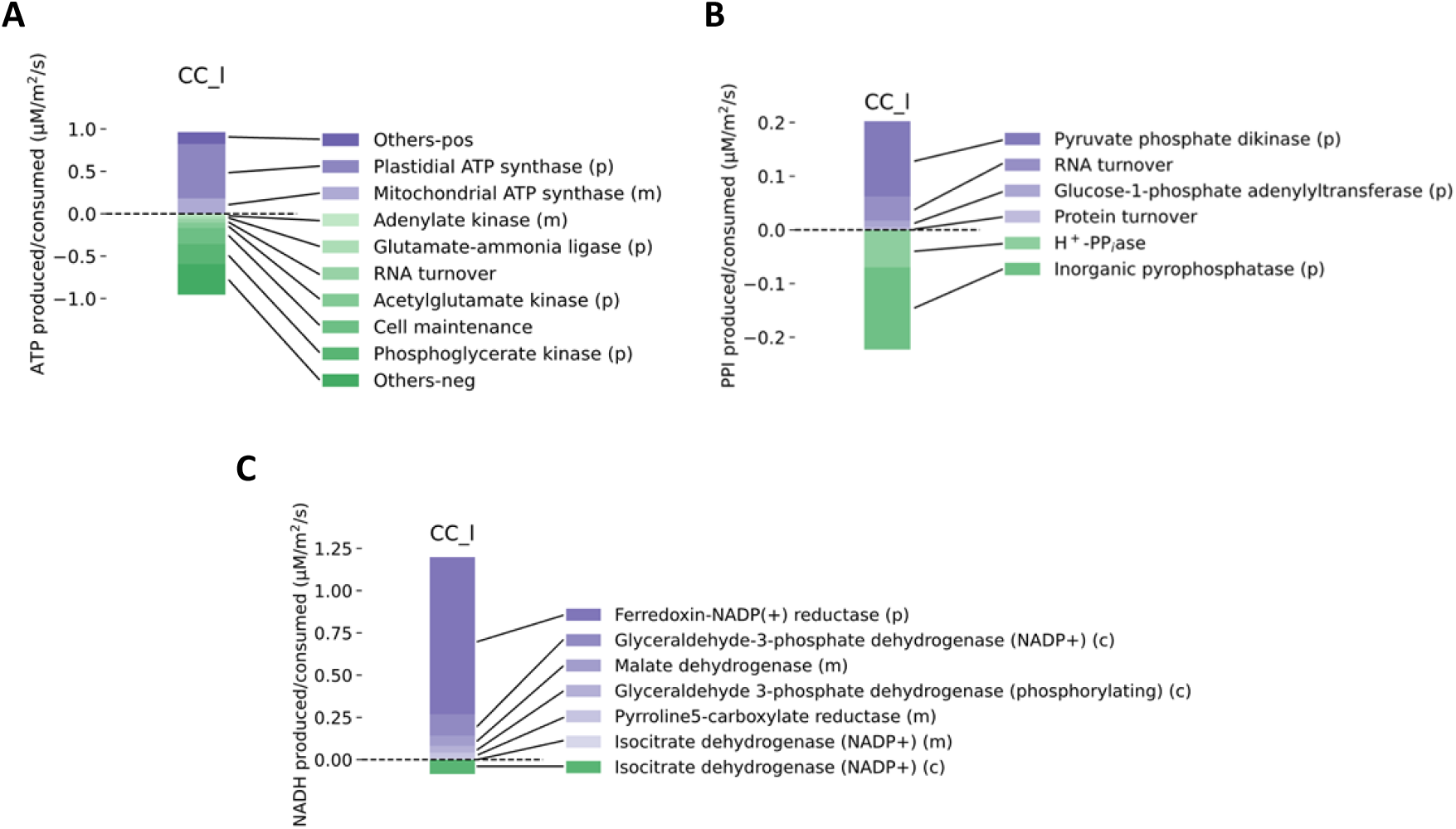
Synthesis and expenditure of energy and reducing power in companion cells during the light phase. (A) A plot of the major reactions involved in ATP synthesis and consumption in companion cells during the light phase. (B) A plot of the major reactions involved in PPI synthesis and consumption in companion cells during the light phase. (C) A plot of the major reactions involved in NAD(P)H synthesis and consumption in companion cells during the light phase. (m, (p), and (c) indicate reaction occurs in the mitochondrion, chloroplast, and cytosol respectively.

### H^+^-PP_i_ase as a pyrophosphate synthase

It has been proposed that the H^+^-PP_i_ase could work in reverse, exploiting the proton gradient either at the plasma membrane or tonoplast to allow pyrophosphate synthesis. Photoassimilate loading into companion cells is an energy intensive process and sucrose is necessarily highly concentrated in companion cells. Catabolising a portion of the imported sucrose to generate the ATP required to import more of it is an effective solution to the high ATP cost of photoassimilate import into the phloem. In the sucrose catabolism pathway, several steps, namely the conversion of UDP-glucose to glucose-1P and the phosphorylation of fructose-1P to fructose 1,6-bisphosphate, can be powered through either ATP or PP_i_ hydrolysis. Higher PP_i_ concentrations in the cytoplasm could increase the rate of these reactions, increasing the rate of sucrose catabolism and production of the ATP generated (Gaxiola et al. 2012, 2016; Primo et al. 2019; Schilling et al. 2017).

Additionally, it has been calculated that it is thermodynamically unlikely for a vacuolar H^+^-PP_i_ase protein to be hydrolysing PP_i_ at 20° C given available data on cytosolic ion concentrations and pH (Davies 1997; Davies et al. 1993). As the membrane potential difference is higher at the plasma membrane, the same arguments apply. While there is evidence of H^+^-PP_i_ase acting in both the PP_i_ hydrolysing and synthesising directions, its function at the plasmalemma of companion cells has yet to be unequivocally determined. It is quite possible that it is situation specific (Bao et al. 2009; Rocha Façanha and de Meis 1998; Seufferheld et al. 2004).

We do not see any PP_i_ synthesis via H^+^-PP_i_ase in any of our model solutions. In fact, due to the amount of PP_i_ produced from biosynthesis, our transcriptome-weighted phloem model indicates that H^+^-PP_i_ase is the dominant proton pump in *Arabidopsis* companion cells. Further, flux variability analysis indicates that the pump running in reverse, to synthesise PP_i_, would reduce phloem export (Supplemental Table ST1). This prediction can be traced to the constraint on carbon intake in mesophyll cells which limits the amount of sucrose they can produce and that companion cells can then import. Companion cells have smaller chloroplasts and the added ATP drain of importing photoassimilates so there is an energy bottleneck at the companion cell-apoplast interface.

While our model indicates that PP_i_ synthesis via H^+^-PP_i_ase would be counterproductive in maximising phloem output, complete reliance on H^+^-PP_i_ase as predicted by our model is contradicted by the results of H^+^-ATPase knockout (Haruta et al. 2010; Zhao et al. 2000). One explanation for this, is that the in *vivo* capacity of the H+-PPiase proton pump is considerably lower than specified in our model where the upper bounds for flux through both H^+^-ATPase and H^+^-PP_i_ase are due solely to cell ratio constraints and do not take into account the actual abundance and kinetics of these proteins. It is likely that the abundance of both proton pumps is a limiting factor in the flux through these reactions (Khadilkar et al. 2016; Robertson et al. 2004). To explore this further in silico, we re-ran the model limiting H^+^-PP_i_ase to 10% of its previous flux. This results in the H^+^-ATPase pump taking over as the dominant plasmalemmal proton pump in companion cells and an increase in the flux through cytosolic inorganic pyrophosphatase (Supplemental Table ST3). Notably, it also reduces the rate of phloem transport by approximately 3%, confirming that sole utilisation of H^+^-PP_i_ase to energise the companion cell plasma membrane is a stoichiometrically more efficient solution. Furthermore, when we investigated an optimised model (parsimonious FBA) without any transcriptome weighting, H^+^-PP_i_ase was still the preferred pump due to the pyrophosphate synthesised in protein and mRNA turnover (Supplemental Table ST1, Supplementary Figure 2). Additionally, pyrophosphate production requires additional enzymes to catalyse the necessary reactions and these consume ATP which would otherwise directly power the H^+^-ATPase pump. Flux variability analysis of the transcriptome-weighted model solution indicated that there are several viable pyrophosphate producing pathways such as those that involving GDP-glucose pyrophosphorylase, and pyruvate orthophosphate dikinase. Many of these pathways are also involved in polysaccharide synthesis and the transcriptome abundance of the encoded enzymes in several of them suggests there may be some companion cell polysaccharide synthesis, such as in cell wall maintenance (Wolf et al. 2012) and callose formation in sieve elements. There is evidence of dynamic callose formation and degradation at phloem plasmodesmata and, potentially sieve plates, to control flow (Barratt et al. 2011; Vatén et al. 2011; Xie and Hong 2011). The production of either of these carbohydrates would result in a net conversion of ATP to PP_i_ which can then also be used to pump protons out of the cell.

### Transcriptome-weighted model predictions of metabolic activity in sieve elements predict surprisingly high rates of fructose catabolism

According to our model 54% of the sucrose imported into the phloem tissue (note that 70% of mesophyll-fixed carbon is imported into phloem, the discrepancy in the two numbers being due to the uptake of amino acids as well as sucrose into the phloem system) is partially consumed in the sieve elements to meet maintenance costs. Surprisingly, a similar amount of fructose (44% of the total amount of sucrose imported) is predicted to be imported for catabolism within phloem cells (Figure 4). In addition, aspartate, glutamate, proline, and serine are also predicted to be catabolised in sieve elements to produce ATP or reducing power and other amino acids.

These predictions are consistent with the high transcript abundance of fructokinase and enzymes of glycolysis in phloem tissue. Clément et al. 2018 detected relatively high levels of fructose in both the laminae and midveins of *Brassica napus* leaves with higher concentrations in the laminae. Fructose is a plausible source of reducing power for phloem tissue despite the ATP requirements for its catabolism (Figure 2) and provides the largest fractional contribution to the reducing power in petiolar sieve elements in our model. Additionally, several hexose symporter proteins are expressed equally in both the mesophyll and companion cell transcriptomes (Kim et al. 2021) however the high fructose uptake, as well as the dark phase glucose uptake by companion cells is unexpected. Removing the hexose import reactions from our companion cell model resulted in an increase in sucrose import corresponding to the fructose import flux in both the light and dark phases and a decrease in overall phloem export indicating that it is more efficient to import hexoses into the phloem than to break down imported sucrose (Supplementary Table ST4).

Our assumption that sieve element maintenance and protein turnover costs would be similar to those of mesophyll cells on a per cell basis is likely to be an overestimate, especially given their relative lack of organelles and protein synthesising machinery. Despite this, our model analysis indicates that the phloem output of the system is maximised when the sieve elements are able to catabolise sugars present in phloem sap and fulfil the energy costs associated with keeping themselves alive and intact. Given the role of mitochondria in this energetic metabolism, it is relevant to consider that sieve element mitochondria are more rudimentary than those found in other cell types (Behnke et al. 1990; Liu et al. 2022). However, the available evidence suggests that they are still present and metabolically active in mature sieve elements (Esau and Cronshaw 1968; Lee et al. 1971; McGivern 1957; Moninger et al. 1993) and this indicates that they do perform some function within these cells. Whether they are capable of providing the energy needs of sieve elements and how much energy sieve elements actually require to meet their maintenance costs is not known. Halving the sieve element maintenance costs in the model reduces the load on the sieve element mitochondria, reducing the both the sucrose synthase and invertase activity within them, and almost doubles the total phloem output of the model (Supplementary Table ST5) indicating a potentially significant drain on plant resources depending on their metabolic activity which at a minimum must be enough to keep them alive and sustain active retrieval of leaked photoassimilates.

Proteomic surveys of phloem sap have found measurable amounts of proteins belonging to both amino acid synthesis and carbohydrate catabolism (Batailler et al. 2012; Carella et al. 2016; Liu et al. 2022). However, a recent proteome analysis of sieve elements in *N. tabacum* showed a lack of key protein subunits necessary for several amino acid synthesis reactions (Supplemental Table ST6). Removal of these reactions from sieve elements in our model shifts arginine, lysine, and valine biosynthesis from sieve elements to being solely synthesised within mesophyll cells. Additionally, approximately 10% of model reactions could not be matched to protein fragments in the Liu et al. 2022 dataset. Removal of these from modelled sieve elements further reduces the total phloem output of the model. Otherwise, the model remained largely unchanged (Supplemental Table ST7).

### Comparison between model solution and Kim et al. interpretation

While not all metabolic pathways identified in the transcriptome dataset were present in our model, there was general agreement between our model predictions and those made solely from the transcriptome data (Kim et al., 2021) with the exception of amino acid synthesis and degradation pathways (Supplementary Table 2). Kim et al., 2021 predicted greater biosynthesis of glutamine and arginine in phloem cells but lower biosynthesis of leucine, methionine, lysine, valine, glycine, isoleucine, histidine, and phenylalanine in phloem cells compared to the average leaf cell. We see greater biosynthesis of glutamine and arginine in phloem cells in our model on a per cell basis but our model also predicts higher biosynthesis of lysine and histidine for the reasons we mentioned earlier (Figure 5). This difference could be because of an overestimation of companion cell and sieve element metabolic capacity in our model but is also a plausible method of transferring reducing power to phloem cells. Of the other pathways predicted by Kim et al. to have higher activity in companion cells, most of them have higher activity in the dark phase but lower activity in the light phase in our model solution. During photosynthesis (in the light phase) the metabolic flux of most active pathways is elevated in mesophyll cells however at night in many of these pathways (non oxidative PPP, glycolysis, TCA cycle, aerobic respiration) companion cells are much more active than mesophyll cells. The high transcriptome abundance for enzymes in these pathways could potentially be a result of the time of measurement or an indication of the higher enzyme concentration requirements to ensure adequate reaction flux in the dark phase.

### Flexible distribution of reactions between cells of the phloem

Due to the free transport of metabolites via the symplast between companion cells and sieve elements in our model, solutions in which a reaction occurs in the companion cells are equivalent in ‘cost’ to solutions in which the same reaction occurs in the sieve elements instead. For example, the pyrophosphate generating loop depicted in Figure 2 could happen almost entirely within companion cells with no effect on modelled phloem output. However, maintenance costs in the sieve elements require that at least some ATP and NADPH generating reactions must occur within them. It is due to this that the model solution suggests that the most efficient cells to produce the histidine, leucine, tyrosine, and valine required for phloem output are sieve elements as they are produced from products of the pentose phosphate pathway and glycolysis. The source of ATP and reducing power in sieve elements is likely to correlate highly with where these amino acids are synthesised and evidence of either happening in sieve elements would tell us a lot more about the extent of sieve element metabolism. Whether these patterns of metabolite diffusion between the symplasts of phloem cells are achievable would depend on enzyme kinetic and regulatory considerations in order to establish the relevant metabolite concentration gradients, something that is beyond the scope of this work.

### Future Perspectives

While FBA models are capable of producing remarkably accurate flux predictions (Cheung et al. 2014; Williams et al. 2010), they are necessarily simplified approximations of the system we are modelling. More data would allow us to make further refinements to our model. Improvements such as higher resolution of the time dependent changes to the cell metabolism (Töpfer et al. 2020), ideally with transcript data from different times of day, or greater thermodynamic constraints based on metabolite concentrations might reveal further aspects of phloem metabolism that we have overlooked. Additional cell-type specific data, especially from proteomic and metabolomic analysis would allow a more detailed assessment of the distribution of metabolism between the individual cell types.

To test the predictions made here, further investigation into the capabilities of companion cell chloroplasts would be necessary. For example, genetic interventions such as knockout of RuBisCO in a companion-cell-specific manner. Alternatively, measurements of the metabolite gradients between companion cells and adjacent sieve elements would give us a better idea of the likely directions of symplastic transport and ATP/ADP in companion cell chloroplasts and cytosol under varying light intensities and/or with varying access to sucrose to test the energetic link between them may also point towards the major source of companion cell ATP (De Col et al., 2017).

Overall, we have clarified several unknowns in phloem metabolism and our work may be the basis for further work to improve the efficiency of phloem loading and hence plant growth rate. This could ultimately be important both in mitigating against rising CO_2_ levels and enhancing crop productivity.

## Methods

Each cell within our model contains a stoichiometric model of core cell metabolism based on the model described in (Shameer et al. 2019). The core model is available in SBML at (https://github.com/hilaryh/phloem)). The metabolic network present within each cell is nearly identical. The only differences are: lack of a vacuole and mRNA turnover reaction in sieve elements; only mesophyll cells and companion cells can exchange sugars and amino acids with the apoplast; companion cells have a H^+^-pyrophosphate pump (AVP1) on their plasma membrane which allows them to break down pyrophosphate to export protons or, conversely, import protons to synthesise pyrophosphate; and sieve elements in the petiole have an additional phloem export reaction that fixes the ratios of amino acids and sucrose.

### Comparative populations of each cell type

The ratio of mesophyll cells to vascular cells in mature Arabidopsis leaves is approximately 1 : 1 (Pyke et al. 1991). Leaf vascular tissue is composed of xylem, phloem, cambium, and the bundle sheath. Phloem accounts for approximately 30% of the vascular tissue in Arabidopsis stems (Collins et al. 2015). Assuming this ratio within vascular tissue remains the same in leaves, and that the phloem is composed of sieve elements and companion cells in the ratio is 5 : 1, the ratio of mesophyll cells to sieve elements to companion cells is 20 : 5 : 1 in leaves.

Sieve elements are approximately 100 μm long and there are approximately 100 in the cross section of a petiole. In a petiole of length 15 mm, we would therefore expect ∼ 15000 sieve elements. There are ∼50 000 mesophyll cells in an Arabidopsis leaf (Pyke et al. 1991) so the ratio of mesophyll cells to companion cells to sieve elements to petiolar sieve elements is approximately 20:1:5:6. To prevent modelled phloem cells undertaking an implausibly high proportion of leaf tissue metabolism, the absolute values of the upper and lower bounds of each reaction in companion cells and sieve elements were therefore constrained to be less than or equal to the flux through the most active mesophyll cell reaction (usually Photon_ep). Further, the bounds on maintenance fluxes, including protein and mRNA turnover, and carbon uptake constraints were additionally scaled by cell ratios. For example, any reactions with an upper bound of 20 μM/m^2^/s in mesophyll cells had an upper bound of only 1 μM/m^2^/s, 5 μM/m^2^/s and 6 μM/m^2^/s in companion cells, sieve elements, and petiole sieve elements respectively.

### Companion cell chloroplasts

While the influence of chloroplast size on photosynthetic efficiency appears to be complicated (Li et al. 2013; Xiong et al. 2017), companion cells have fewer and smaller chloroplasts than mesophyll cells and there are approximately on 5% as many companion cells as mesophyll cells in the leaf. So we have estimated that companion cell chloroplasts can capture no more than 1% of the photons captured by mesophyll chloroplasts and constrained the companion cell photon uptake reaction (Photon_ep_CC_l) accordingly.

### Protein/RNA turnover

Protein degradation rates in mature Arabidopsis leaves are highly variable but the degradation rates, K_d_, of known metabolic proteins tend to magnitudes of approximately 0.1 per day (Li et al., 2017) or approximately 10^−6^ per second. The fold change in protein, ΔP, over time can be calculated as

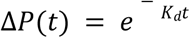

ΔP is then 10^−6^ per second. There are approximately 15 mg protein per gFW in Arabidopsis leaves (Piques et al., 2009). Assuming the average molecular weight of each amino acid is 118.9 g/mol (Hachiya et al., 2007), we arrive at an amino acid density of 0.13 mmol AA/gFW. With a FW to DW ration of approximately 50:1, we estimate the DW protein density to be 6.3 mmol AA/gDW.

The fold change in amino acids is then 6.3 ∗10^−9^ mol AA/gDW/s. The maximum estimated rate of protein synthesis is 26 µmol AA /gFW/day, about twice our estimate. Leaf mass per area is approximately 1.5 mg DW/cm^2^ = 15 gDW/m^2^ (Liu et al., 2020). Then the flux through our protein turnover reaction is 9.45∗10^−2^ µmol AA/m^2^/s. The total RNA present in Arabidopsis cells varies throughout the day. On average, it is approximately 600 ug/gFW (Piques et al., 2009). The half-life of mRNA is approximately 107 mins (Sorenson et al., 2018; Szabo et al., 2020). Thus, the fold change in mRNA is 10^−4^ s^-1^.

The average molecular weight of a ribonucleotide is 500 Da. Then the flux through our RNA turnover reactions should be 0.9 µmol/m^2^/s.

### Weighting FBA model with transcriptome data

We used cell-specific transcriptome data published by (Kim et al. 2021) to weight model solutions towards those more likely to represent *in vivo* cell metabolism using a modification of the RegrEx method developed by (Robaina Estévez and Nikoloski 2014; Robaina-Estévez et al. 2017). For companion cell and sieve element reactions where companion cells synthesise the transcripts for both cell types, the objective was to minimise the difference between the sum of fluxes and their corresponding transcript abundance in companion cells. The objective can be written as:

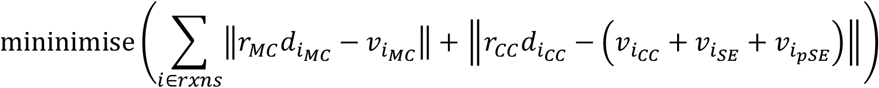

where *r*_*cell*_ is the ratio of the cell type in leaves (i.e. *r*_*MC*_ = 20, *r*_*CC*_ = 1), *d*_*icell*_ is the transcript abundance of reaction *i* in *cell*, and *v*_*icell*_ is the flux through reaction *i* in *cell*. Instead of the LASSO method used to determine the relative weights of transcriptome data and sum of fluxes, we prioritised the similarity to transcriptome data and then identified a solution with the minimum sum of fluxes (parsimonious) within that solution space.

To check the stability of our observations, flux variability analysis (FVA) (Mahadevan and Schilling 2003) was performed on both the transcriptome-weighted and parsimonious versions of our model. The FBA solutions were generated using gurobipy and the COBRAPy package (Ebrahim et al. 2013). Python code required to reproduce the results in this article can be found at (https://github.com/hilaryh/phloem).

## Supplementary Information

The following supplemental materials are available:

**Supplemental Figure S1** A flux diagram of mesophyll cell metabolism in the RegrEx model solution

**Supplemental Figure S2** A flux diagram of companion cell metabolism in the parsimonious model solution

**Supplemental Figure S3** ATP budgets for each cell in the RegrEx model solution

**Supplemental Figure S4** PPi budgets for each cell in the RegrEx model solution

**Supplemental Figure S5** NADH budgets for each cell in the RegrEx model solution

**Supplemental Table ST1** Model reactions, their stoichiometries, and fluxes in the parsimonious and RegrEx model solutions as well as flux variability analysis ranges. Fluxes for model solutions without PSII in companion cells, and with AVP1 reversed are also provided

**Supplemental Table ST2** Comparison of pathway activity between transcriptome data-based and transcriptome-weighted model predictions

**Supplemental Table ST3** Predicted reaction fluxes when AVP1 activity is restricted to only 10% of the optimal

**Supplemental Table ST4** Predicted reaction fluxes when hexose uptake is prevented in model companion cells

**Supplemental Table ST5** Predicted reaction fluxes when sieve element maintenance costs are halved

**Supplemental Table ST6** Mapping of *N. tabacum* sieve element proteome data onto model reactions

**Supplemental Table ST7** Predicted reaction fluxes when proteins not found in *N. tabacum* dataset are removed from modelled sieve elements

## Author contributions

SV, CF, ARF, US, and LJS conceived the study; LJS supervised the research; HH and LJS developed the modelling approach; HH implemented the model; all authors provided critical feedback and contributed to writing the manuscript.

## Acknowledgements

We thank Sanu Shameer (University of Oxford) for helpful discussion and Joerg Hofmann (Friedrich-Alexander-University) for help in interpreting the proteome data referenced. We also thank BASF for funding this project.

